# Urbanization-driven changes in web-building are decoupled from body size in an orb-web spider

**DOI:** 10.1101/214924

**Authors:** Maxime Dahirel, Maarten De Cock, Pieter Vantieghem, Dries Bonte

**Affiliations:** Ghent University, Department of Biology, Terrestrial Ecology unit, K. L. Ledeganckstraat 35, B-9000 Gent, Belgium; Univ Rennes, CNRS, ECOBIO (Ecosystèmes, biodiversité, évolution) - UMR 6553, F-35000 Rennes, France

**Keywords:** adaptation, fecundity, foraging, multivariate mixed model, spider web, temperature-size rule

## Abstract

1. In animals, behavioural responses may play an important role in determining population persistence in the face of environmental changes. Body size is a key trait central to many life history traits and behaviours. While behaviours are typically assumed to be highly plastic, size correlations may impose constraints on their adaptive value when size itself is subject to environmental changes.
2. Urbanization is an important human-induced rapid environmental change that imposes multiple selection pressures on both body size and (size-constrained) behaviour. How these combine to shape behavioural responses of urban-dwelling species is unclear.
3. Using web-building, an easily quantifiable behaviour linked to body size, and the garden spider *Araneus diadematus* as a model, we disentangle direct behavioural responses to urbanization and body size constraints across a network of 63 selected populations differing in urbanization intensity at two spatial scales.
4. Spiders were smaller in highly urbanized sites (local scale only), in line with expectations based on reduced prey biomass availability and the Urban Heat Island effect. The use of multivariate mixed modelling reveals that although web traits and body size are correlated within populations, behavioural responses to urbanization do not appear to be constrained by size: there is no evidence of size-web correlations among populations or among landscapes. Spiders thus altered different components of their web-building behaviours independently in response to urbanization: mesh width and web surface decreased independently with urbanization at the local scale, while web surface also increased with urbanization at the landscape scale. These responses are expected to compensate, at least in part, for reduced prey biomass availability.
5. Our results demonstrate that responses in typically size-dependent behaviours may be decoupled from size changes, thereby allowing fitness maximisation in novel environments. The spatial scale of the behavioural responses to urbanization suggest contributions of both genetic adaptation and plasticity. Although fecundity decreased with local-scale urbanization, *Araneus diadematus* abundances were remarkably similar across urbanization gradients; behavioural responses thus appear overall successful at the population level.

## Introduction

In animals, behaviour is often considered as the first route of adaptation to rapid environmental changes (Wong & Candolin, 2015). Numerous examples of both adaptive and maladaptive behavioural changes in response to human-induced environmental changes have now been recorded (Lowry, Lill, & Wong, 2013); the potential costs and constraints associated with these behavioural changes are, however, poorly understood. Behaviours may be linked to metabolic and physiological processes that directly impact fitness (Bonte et al., 2012; Debecker, Sanmartín-Villar, de Guinea-Luengo, Cordero-Rivera, & Stoks, 2016). When behaviour is correlated to other traits, conflicting selection pressures may hinder adaptation, leading to mismatches between the expressed and optimal behaviours in the new environment (Sih, 2013; Wong & Candolin, 2015). In particular, many behaviours are correlated to body size (e.g. Gregorič, Kuntner, & Blackledge, 2015; Stevens et al., 2014), a key trait that can itself be directly impacted by environmental changes (Oliveira, Freitas, Scheper, & Kleijn, 2016; Renauld, Hutchinson, Loeb, Poveda, & Connelly, 2016).

Urbanization is one of the most prominent human-induced environmental changes, with cities now harbouring more than half of the global human population (Seto, Güneralp, & Hutyra, 2012; United Nations Population Division, 2015). Direct and indirect impacts of the urbanization process include habitat fragmentation, increased temperatures (the “Urban Heat Island” effect), elevated levels of pollution and changes in resource availability due to the decline of key species and/or the increased availability of anthropogenic food sources (Alberti, 2015; Parris, 2016). This cocktail of environmental changes means cities present novel ecological conditions never encountered before in organisms’ evolutionary histories (Alberti, 2015; Hendry, Gotanda, & Svensson, 2017; Johnson & Munshi-South, 2017). This drives changes in community taxonomic and functional composition (Dahirel, Dierick, De Cock, & Bonte, 2017; Piano et al., 2017) as well as intraspecific phenotypic changes (plastic and/or genetic: Alberti et al., 2017; Brans, Jansen, et al., 2017; Johnson & Munshi-South, 2017; Lowry et al., 2013) at an unprecedented rate (Alberti et al., 2017). Cities can therefore be seen as key “natural experiments” to understand eco-evolutionary consequences of global change (Johnson & Munshi-South, 2017).

Orb web spiders (Araneae, Araneidae) are a unique model for the study of behaviour, as their foraging behaviour is archived in their web (Foelix, 2010; Fig. 1). Detailed quantification of the orb web structure therefore allows inferences on both the implemented foraging strategies (from web architecture) and the energetic investments in web production (Sherman, 1994). Keeping total investment equal, larger spacing of the capture threads allows the construction of larger webs and an increase in prey interception; conversely, pressures to build small meshes to e.g. increase prey retention (Blackledge & Zevenbergen, 2006) come at the cost of reduced capture areas (Sandoval, 1994). Spiders can maximise both retention and interception simultaneously only by increasing the quantity of capture silk per web (Eberhard, 2013; Sherman, 1994). Variation in web building behaviour has been recorded in relation to aging and growth (e.g. Witt, Rawlings, & Reed, 1972), and in response to a wide variety of environmental changes. Spiders experiencing lower prey biomass have for instance been shown to increase the capture area of their web to increase its efficiency (Mayntz, Toft, & Vollrath, 2009). Spiders may also use information on the size of recently caught prey to optimize the web mesh width and capture area (Schneider & Vollrath, 1998). The architecture and size of webs is influenced by weather, both through changes in the physical pressures the web has to withstand and (especially temperature) through changes in the internal physiological state of the building spider (Barghusen, Claussen, Anderson, & Bailer, 1997; Vollrath, Downes, & Krackow, 1997). Web building can also be altered by exposure to various chemical pollutants (Benamú, Schneider, González, & Sánchez, 2013), and constrained by space availability in the current habitat (Vollrath et al., 1997). Most of these environmental drivers are highly altered by urbanization (Dahirel et al., 2017; Parris, 2016) and may either impose constraining or adaptive changes in web building.

**Figure 1.**
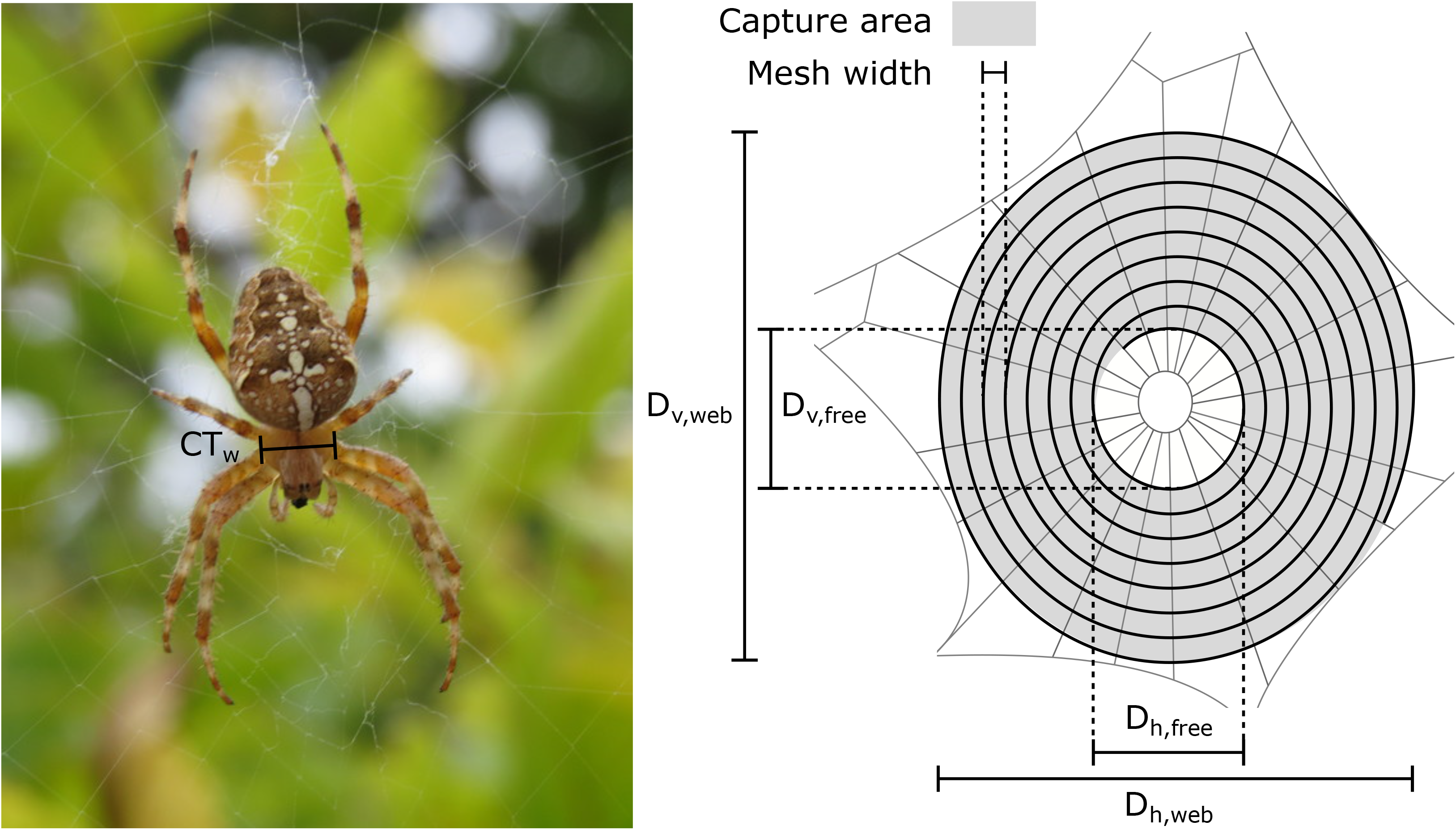
Left: photograph of a female *Araneus diadematus* showing the distinctive cross pattern on the abdomen (credit: Maxime Dahirel); CT_w_: cephalothorax width, used as a proxy of body size. Right: diagram of an *Araneus diadematus* web, with radii spanning from the central hub supporting a sticky silk thread spiral (bold). D_v_ and D_i_ refer to the vertical and horizontal diameters of the full web (D_web_) and the free zone (D_free_), used to estimate the capture area (in grey); the number of threads of the capture spiral on the horizontal and vertical axes of the capture area is used to calculate average mesh width.

Besides these direct responses, environmental changes may also influence spider foraging behaviour indirectly, as web-building is coupled to body size both within and among species. Larger spiders produce on average larger, and often less densely structured webs (Gregorič et al., 2015; Heiling & Herberstein, 1998). As increased temperatures in urban areas are expected to lead to reduced body size in ectotherms through increases in metabolic rates (Brans, Jansen, et al., 2017; Horne, Hirst, & Atkinson, 2015; Sheridan & Bickford, 2011), this may for instance lead to additive reduced investments in web-building. These size-dependent effects may be compounded when urbanization influences resource availability during development (Dahirel et al., 2017), as spider adult size is also resource-dependent (Mayntz, Toft, & Vollrath, 2003). Opposing selection pressures related to urbanization may thus constrain optimal web-building behaviour. Furthermore, since web-building behaviour is energetically costly (Blackledge, Kuntner, & Agnarsson, 2011), it is directly dependent on the total availability of energy. One can therefore expect the maintenance of optimal web-building strategies to be traded-off with other life histories. By relating web-building behaviour, individual size and fecundity, it should thus be possible to understand how changes in environmental conditions directly, but also indirectly affect the adaptive value of these behavioural changes.

Behavioural responses to urbanization and other human-induced rapid environmental changes, whether adaptive or not, can result from plasticity and/or genetic changes (Sih, Stamps, Yang, McElreath, & Ramenofsky, 2010; Tuomainen & Candolin, 2011). While a formal understanding of the underlying sources of phenotypic (behavioural) variation in response to urbanization can only be achieved by experimental work (Johnson & Munshi-South, 2017; Merilä & Hendry, 2014), the spatial scale of phenotypic variation, relative to the scale of dispersal, may provide first insights. Spiders disperse by ballooning using silk threads as a sail for airborne travel (Bonte, 2012); although very long-distance dispersal events occur, most dispersers are thought to land within a few hundred metres from their departure point (Bell, Bohan, Shaw, & Weyman, 2005; Reynolds, Bohan, & Bell, 2007). Once airborne, individuals have no control over their trajectory, meaning adaptive habitat choice (Edelaar & Bolnick, 2012) is unlikely. Gene flow is thus expected to swamp genetic differentiation at these spatial scales, and any response to environmental variation with a spatial grain equal or smaller than the typical dispersal event more likely to result from plasticity. By contrast, evidence of phenotype-environment matching at larger spatial scales (several kilometres) suggests putative genetic adaptation (Macdonald, Llewelyn, & Phillips, 2018; Richardson, Urban, Bolnick, & Skelly, 2014).

Here, we study and quantify shifts in body size and web building in response to urbanization, using univariate and multivariate mixed models in order to identify the relative contribution of size constraints to (adaptive) behavioural changes. We used the garden spider *Araneus diadematus* as a model; this species is one of the most common species in both urban and non-urban orb web spider communities in western Europe (Dahirel et al., 2017). *A. diadematus* alters its web-building behaviour depending on abiotic conditions and the availability/ characteristics of potential prey (Bonte, Lanckacker, Wiersma, & Lens, 2008; Schneider & Vollrath, 1998; Vollrath et al., 1997); webs are recycled and rebuilt daily, allowing spiders to match currently/recently experienced environmental conditions (Breed, Levine, Peakall, & Witt, 1964). We used a well-studied network of urban and non-urban sites in which differences in resource availability and temperature have been previously recorded (Dahirel et al., 2017; Kaiser, Merckx, & Van Dyck, 2016; Merckx et al., 2018). Although prey numbers are roughly constant across urbanization levels, prey biomass is expected to be lower in highly urbanized sites as the size of the average prey declines by about 18% (Dahirel et al., 2017). Average temperatures are higher in highly urbanized sites, but this effect is mainly detected when considering urbanization at the most local scale (about +1-2°C) and weaker to non-significant at larger spatial scales (Merckx et al., 2018). We therefore expected *Araneus diadematus* to present adaptive shifts in web-building behaviour in response to urbanization, despite reduced size (due to reduced resources and/or the Urban Heat Island effect) and body size-related constraints expected to lead to reduced web production in cities. We studied responses to urbanization at two independent spatial scales, to obtain indices on the roles of genetic adaptation vs. plasticity, and to account for the fact that environmental correlates of urbanization may be scale-dependent (Kaiser et al., 2016; McDonnell & Hahs, 2015; Merckx et al., 2018). We additionally analyse spider abundance and fecundity data to investigate potential fitness consequences of adaptation to city life.

## Material and Methods

### Study species

*Araneus diadematus* Clerck 1757 is a common orb-weaving spider present across the Holarctic in a wide range of natural and human-altered environments, building its web in shrubs and tall herbaceous vegetation (Lee & Thomas, 2002; Nentwig, Blick, Gloor, Hänggi, & Kropf, 2016). Its distinctive dorsal cross pattern makes field identification easy (Roberts, 1993; Fig. 1). Females usually become mature in late summer, and can survive through to late autumn (Lee & Thomas, 2002). In cities, *Araneus diadematus* mostly settles in gardens and greenspots as opposed to roadsides or close to buildings (Van Keer, Vanuytven, De Koninck, & Van Keer, 2010).

### Study sites

We sampled 63 *A. diadematus* populations across a well-studied network of urban, rural and natural landscapes in northern Belgium (Brans, Govaert, et al., 2017; Dahirel et al., 2017; Kaiser et al., 2016; Merckx et al., 2018; Piano et al., 2017; Supplementary Figure S1), one of the most urbanized and densely populated regions in Europe (United Nations Population Division, 2015). Urbanization was studied at two different spatial scales thanks to a 2-step stratified selection design, with 3 × 3 km landscapes (“landscape scale” urbanization) being selected first, then three 200 × 200 m sites (“local scale” urbanization) chosen within each landscape. We used the percentage of surfaces occupied by buildings (extracted from the Large-scale Reference Database, a reference map of Flanders; https://www.agiv.be/international/en/products/grb-en) as our proxy for urbanization. This metric is precise down to the individual building, and thus usable at both spatial scales, but excludes other artificialized surfaces such as roads, railways, parking spaces or pavements; hence percentages higher than 10% already correspond to highly urbanized contexts. We cross-checked this metric with the percentage of artificial surfaces based on the CORINE land cover database (European Environmental Agency, 2016)(Level 1: “Artificial surfaces”, excluding classes 1.41 and 1.42 corresponding to urban greenspots). This includes all artificialized surfaces but, due to its coarser resolution (minimum mapping unit 25 ha) cannot be used at smaller spatial scales. Both metrics were highly correlated (*N* = 21 landscapes, Spearman correlation coefficient = 0.94, *p* = 5.17 × 10^-6^, Supplementary Figure S2). Twenty-one 3 × 3 km landscapes were selected and sorted into three urbanization levels (7 landscapes by level). High-urbanization landscapes had more than 10% of their area covered by buildings (equivalent to more than 70% of artificial surfaces based on CORINE). Low-urbanization landscapes had less than 3% of their surfaces occupied by buildings (less than 20% of artificial surfaces based on CORINE); additionally, they were selected so that more than 20% of their surface was covered by so-called “ecologically valuable areas” (areas with rare, vulnerable or highly diverse vegetation based on the Flanders Biological Valuation Map; Vriens et al., 2011). A third class of “intermediate” landscapes had between 5 and 10% of their surface covered by buildings (between 20 and 70% of artificial surfaces based on CORINE. Within each landscape, three 200 × 200 m sites (one per urbanization level) were selected this time based on the percentage covered by buildings only. Vegetated areas in chosen sites were unforested and dominated by low vegetation (grasslands/lawns with low shrubs and occasional trees in e.g. gardens, parks or hedgerows).

### Spider collection and phenotypic measurements

Populations were sampled from 25 August to 5 October 2014, i.e. during the first part of the reproductive period (Lee & Thomas, 2002). One landscape (3 sites) was visited per day; there was no significant link between landscape-level urbanization and sampling date (ANOVA; *N* = 21 landscapes, *F_2,18_*= 0.009, *p* = 0.991). In each visited site, between 7 and 11 adult females per population (*N_total_*= 621, average ± SD : 9.86 ± 0.74) were sampled on their webs and stored in 70% ethanol; spiders’ cephalothorax width was measured under binocular microscope and used as a proxy for body size (Bonte et al., 2008; Fig. 1). Out of these 621 spiders, 193 individuals caught in the 9 landscapes of the Ghent region (Supplementary Figure S1) were also dissected and the number of mature eggs recorded. As *Araneus diadematus* webs are conspicuous and located in similar habitats (grassland-shrub mosaics taken in the broad sense) independently of urbanization levels (Lee & Thomas, 2002; Van Keer et al., 2010), we were also able to collect reliable population density data (number of adult female spiders observed per 200 × 200 m site in 4.5 person-hours) in the 62 sites (out of 63) sampled in the community-level study by Dahirel *et al*. (2017).

Based on measurements taken in the field (vertical and horizontal diameters of whole web and free central zone, number of sticky silk spirals in each web quadrant, Fig. 1), we estimated two design parameters for the webs belonging to sampled spiders: the web capture area surface (considering orb webs as ellipses, following Herberstein & Tso, 2000), and the mesh width (interval between sticky spirals, averaged over the horizontal and vertical axes).

### Statistical analysis

All analyses were carried out using R, version 3.4 (R Core Team, 2017). Body size, web traits and sampling date values were scaled to unit standard deviation prior to inclusion in models (see Table 1 for descriptive statistics), in order to improve model convergence and make results comparable across parameters and variables.

**Table 1.**
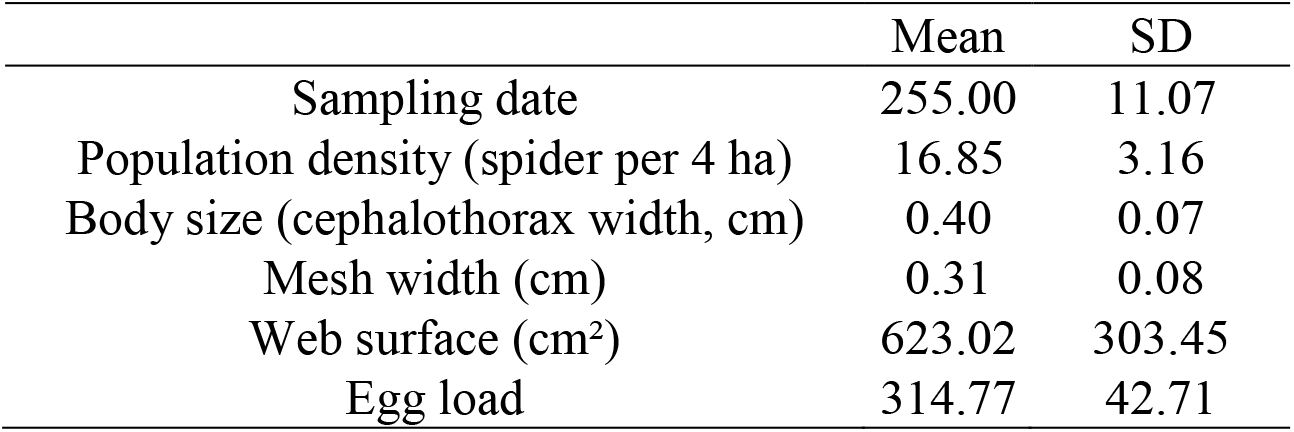
Descriptive statistics for the quantitative traits

The number of adult spiders per site was analysed using a Poisson generalized linear mixed model. Spider numbers were modelled as a function of local and landscape level of urbanization, as well as sampling date, with a random effect of landscape identity added to account for spatial clustering of populations in landscapes.

We follow broadly Araya-Ajoy and Dingemanse (2014)’s approach and analyse spider traits and their correlations in two steps.

First, each trait (body size, web surface, mesh width, egg number) was analysed separately using univariate mixed models. These models included fixed effects of local and landscape level of urbanization as well as sampling date, with random effects of landscape and site of origin to account for non-independence of observations. Body size, web surface and mesh width were analysed using linear mixed models, egg number was analysed using a Poisson generalized linear mixed model.

Second, we used a multivariate mixed model framework (Araya-Ajoy & Dingemanse, 2014; Dingemanse & Dochtermann, 2013) to estimate trait covariances/correlations and partition them across hierarchical levels (correlations among landscapes, among populations, and (residual) within populations). Fitting these covariances is essential to understand whether observed responses to urbanization in one trait are actually due to correlated responses across the different web-building traits and body size. Fixed effects, random intercepts and error families for each trait were included as in univariate models. We fitted two multivariate models differing in their hypothesized covariance structure among the four traits. In model #1, we fitted three covariance matrices, one per hierarchical level (among-landscapes, among-populations, within populations). This assumes there can be non-zero correlations between any pair of trait, and that the strength and direction of these correlations only depend on the scale at which they are considered. In the more complex model #2, we allowed covariance structure to additionally vary depending on urbanization level at the corresponding scale (landscape scale for among-landscapes correlations, local scale for among-populations), leading to 7 covariance matrices being fitted (three per urbanization level at each scale, plus the residual within-population matrix).

All models were fitted using the MCMCglmm R package (Hadfield, 2010). We used uninformative default priors for fixed effects, inverse-Wishart priors for the residual (co)variances and parameter expanded priors for among-landscapes and among-population (co)variance matrices. Degrees of belief ν for the (co)variance matrices were set to weakly informative values, i.e. the dimension of the corresponding matrix (1 for univariate models, 4 for multivariate models). Chains ran for (univariate/ multivariate model #1/ multivariate model #2) 250000/ 2200000/ 10200000 iterations, with the first 50000/ 200000/ 200000 iterations discarded as burn-in and sampling every 100/ 2000/ 10000 iteration post burn-in, resulting in 1000 samples per chain with serial autocorrelation < 0.1 for all model parameters. We ran three chains per model, checked mixing graphically and confirmed chain convergence using the Gelman-Rubin statistic (using the coda R package; Plummer, Best, Cowles, & Vines, 2006; all 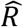 were ≤ 1.05).

There was no evidence of residual spatial autocorrelation in any of the retained models; 95% confidence intervals of spline correlograms (fitted using the ncf package; Bjornstad, 2016) overlapped with 0 at all distances and for all response variables.

## Results

### Spider population density

The number of spiders found per site was not influenced by urbanization at either spatial scale (95% Credible Intervals for all β overlapped with 0, Table 2). There was also no temporal trend in spider abundance (Table 2).

**Table 2.** Sources of variation in population density and spider traits. Parameters from univariate mixed models are presented with 95% Credible Intervals in parentheses. The proportions of among-individual and among-population variation explained by models are presented using Nakagawa and Schielzeth (2013)’s marginal and conditional R² (R²m : variation explained by fixed effects only; R²c: variation explained by fixed and random effects).

|  | Population density (per 4ha) | Spider traits |  |  |  |
| --- | --- | --- | --- | --- | --- |
|  |  | Body size | Mesh width | Web surface | Egg load |
| Fixed effects | $\beta$ (95% CI) | $\beta$ (95% CI) | $\beta$ (95% CI) | $\beta$ (95% CI) | $\beta$ (95% CI) |
| Intercept | 2.70 (2.50; 2.89) | 0.00 (-0.35; 0.35) | 0.24 (-0.13; 0.60) | -0.11 (-0.45; 0.24) | 5.83 (5.77; 5.90) |
| Local urbanization, medium | 0.07 (-0.14; 0.28) | 0.04 (-0.29; 0.38) | -0.14 (-0.39; 0.09) | **-0.25 (-0.47; -0.03)** | **-0.10 (-0.16; -0.05)** |
| Local urbanization, high | 0.04 (-0.17; 0.25) | **-0.46 (-0.81; -0.13)** | **-0.42 (-0.66; -0.18)** | **-0.39 (-0.61; -0.17)** | **-0.18 (-0.23; -0.13)** |
| Landscape urbanization, medium | 0.11 (-0.11; 0.33) | 0.10 (-0.32; 0.52) | -0.13 (-0.60; 0.36) | **0.49 (0.05; 0.93)** | 0.01 (-0.07; 0.08) |
| Landscape urbanization, high | 0.10 (-0.13; 0.32) | 0.32 (-0.11; 0.74) | 0.00 (-0.47; 0.49) | **0.48 (0.01; 0.93)** | -0.03 (-0.10; 0.04) |
| Sampling date | -0.01 (-0.10; 0.09) | 0.14 (-0.03; 0.31) | **0.26 (0.06; 0.45)** | **0.32 (0.13; 0.51)** | 0.07 (-0.00; 0.14) |
| Random effects | $\sigma^2$ (95% CI) | $\sigma^2$ (95% CI) | $\sigma^2$ (95% CI) | $\sigma^2$ (95% CI) | $\sigma^2$ (95% CI) |
| Plot (N=21) | 0.00 (0.00; 0.02) | 0.06 (0.00; 0.21) | 0.15 (0.04; 0.36) | 0.13 (0.04; 0.32) | 0.00 (0.00; 0.00) |
| Site (N=63) | -- | 0.23 (0.12; 0.38) | 0.08 (0.02; 0.17) | 0.06 (0.01; 0.14) | 0.00 (0.00; 0.00) |
| Residual variation | 0.06 (0.03; 0.09) | 0.70 (0.62; 0.79) | 0.74 (0.65; 0.83) | 0.70 (0.62; 0.78) | 0.02 (0.00; 0.02) |
| $R^2_c$ (among individuals) | -- | 0.37 (0.27, 0.48) | 0.33 (0.22, 0.47) | 0.36 (0.25, 0.49) | 0.41 (0.23, 0.63) |
| $R^2_m$ (among individuals) | -- | 0.11 (0.04, 0.20) | 0.12 (0.04, 0.24) | 0.19 (0.08, 0.30) | 0.37 (0.19, 0.59) |
| $R^2_c$ (among populations) | 0.55 (0.43, 0.67) | 0.43 (0.19, 0.68) | 0.77 (0.50, 0.94) | 0.84 (0.64, 0.98) | 0.79 (0.65, 0.91) |
| $R^2_m$ (among populations) | 0.10 (0.02, 0.22) | 0.30 (0.12, 0.49) | 0.37 (0.14, 0.61) | 0.52 (0.26, 0.73) | 0.73 (0.49, 0.88) |

Body size, Mesh width, Web surface and sampling date are centered and scaled to unit standard deviation
Reference category for urbanization levels is "low urbanization"
Family is Gaussian for Body size, Mesh width, Web surface and Poisson for Population density and Egg load
N= 621 except for Egg load (N=193) and Population density (N=63)

### Sources of variation in spider traits: univariate models (Table 2, Fig 2)

Body size, as measured by cephalothorax width, was influenced by the local level of urbanization, with spiders being 8.30% smaller [95% Credible Interval: −14.03%, −2.34%] in highly urbanized sites compared to populations from low-urbanization sites. There was no effect of landscape-scale urbanization or sampling date.

**Figure 2.**
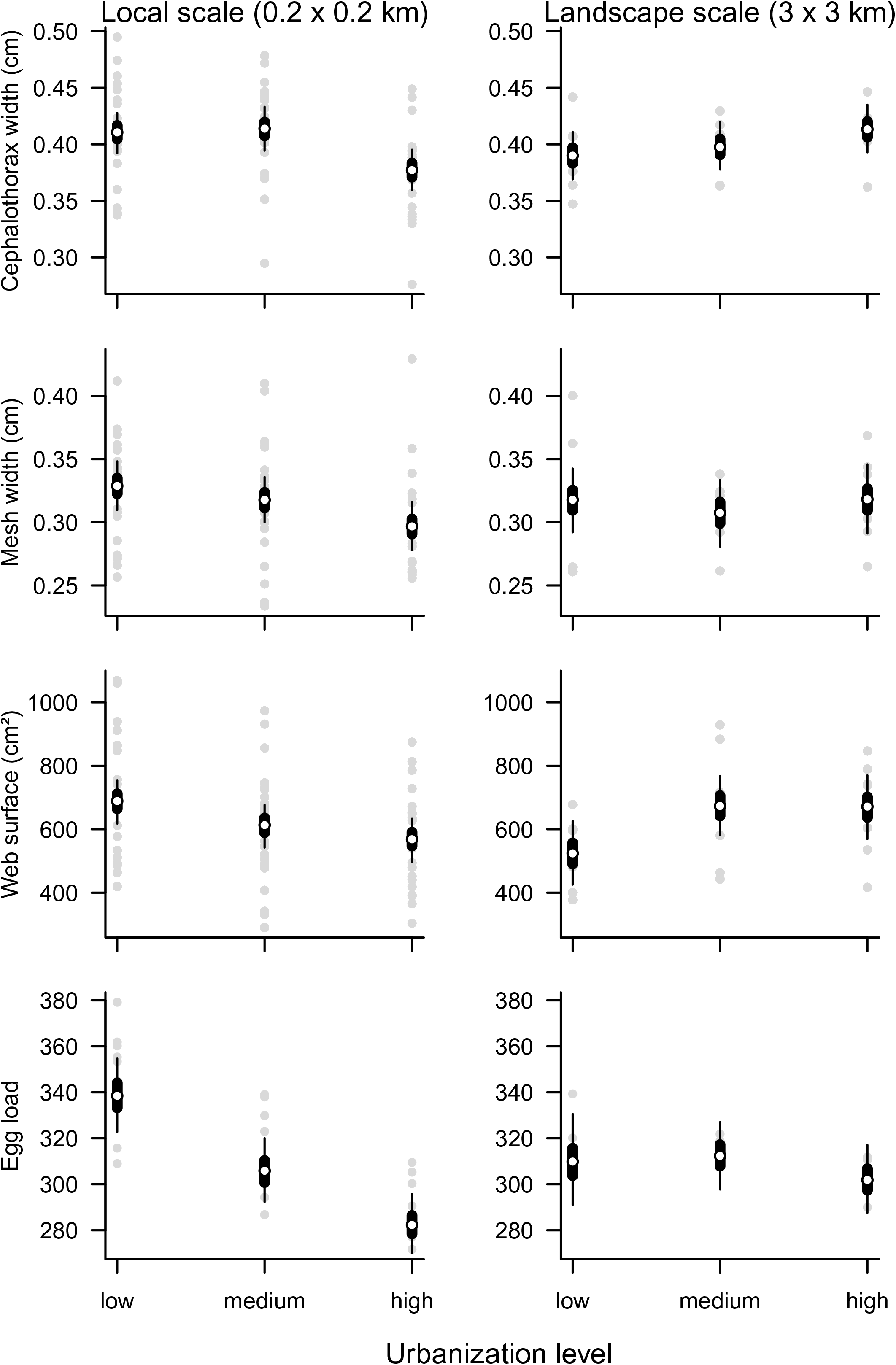
Predicted mean *Araneus diadematus* trait values ± 50% (thick bars) and 95% (thin lines) credible intervals as a function of urbanization intensity at two spatial scales. Predictions are based on fixed effects from univariate models (Table 2), fixed effects from the multivariate model being virtually identical (Supplementary Table S2). Predictions at one spatial scale are made by averaging over the effects of sampling date and urbanization at the other spatial scale. Grey points represent observed individual site (local scale) and landscape average values.

Local urbanization level also influenced mesh width: spiders built webs with narrower mesh in highly urbanized populations compared to low urbanization sites (−9.67% [−14.96%, − 4.07%]). Mesh width was not influenced by landscape-scale urbanization, but increased with sampling date. Web surface decreased with increasing urbanization at the local scale (low to intermediate urbanization: −12.93% [−23.83%, −1.62%]; low to high urbanization: −20.28% [−31.12%, −9.37%]). It increased with landscape-scale urbanization (low to intermediate urbanization: +26.19% [+2.25%, +54.63%]; low to high urbanization: +25.43% [+0.5%, +54.54%]) and with time.

Local urbanization (both intermediate and high levels) decreased fecundity (number of eggs per spider) at the local scale (low to intermediate urbanization: −9.64% [−14.51%, −4.77%]; low to high urbanization: −16.62% [−20.81%, −12.12%]). There was no supported effect of landscape-scale urbanization or sampling date on fecundity.

### Correlations among traits: multivariate models

Although multivariate model #2 allowed for differences in covariance structure due to urbanization, none was found: credible intervals for all random (co)variance estimates largely overlapped across urbanization levels, possibly due to the low number of landscapes/sites per submatrix (Supplementary Tables S1). Further inferences about covariance matrices were therefore made based on the simpler model #1.

Inferences based on fixed effects from this multivariate model were virtually the same as those obtained from the univariate models (see Table 2 vs. Supplementary Table S2 for a comparison of parameters). At the landscape and population levels, none of the estimated trait covariances were different from zero, based on credible intervals (Table 3). At the within-population level, web traits were positively correlated with each other and with body size (Table 3). These results imply that, despite trait correlations existing within local populations, web building traits and body size are jointly but independently diverging in response to local levels of urbanization. To confirm that observed urbanization-induced changes in web surface (Table 2) were independent from the associated changes in mesh width, we refitted this multivariate model with this time no fixed effect of local urbanization for web surface (see Supplementary Tables S3 and S4). Removing this fixed effect did not meaningfully influence the among-population correlation between mesh width and web surface (from 0.02 [−0.02; 0.08] with the main effect to 0.03 [−0.01; 0.11] without, Supplementary Table S4).

**Table 3.** Trait variance-covariance matrices for each hierarchical level under study, based on the retained multivariate mixed model. In each case, among landscapes/populations/individual variances are given on diagonals, with between-trait covariances below and corresponding between-trait correlations above the diagonal. Covariances/correlations involving Egg load are on the latent (log) scale for this trait. 95% credible intervals are given in parentheses

|  | Body size | Mesh width | Web surface | Egg load |
| --- | --- | --- | --- | --- |
| (a) Among landscapes |  |  |  |  |
| Body size | 0.07 (0.00; 0.22) | -0.06 (-0.75; 0.64) | 0.50 (-0.37; 0.92) | -0.05 (-0.89; 0.83) |
| Mesh width | -0.00 (-0.08; 0.08) | 0.17 (0.05; 0.42) | 0.04 (-0.58; 0.58) | -0.02 (-0.78; 0.77) |
| Web surface | 0.05 (-0.01; 0.15) | 0.01 (-0.08; 0.11) | 0.13 (0.03; 0.31) | -0.08 (-0.87; 0.80) |
| Egg load | -0.00 (-0.01; 0.01) | -0.00 (-0.02; 0.02) | -0.00 (-0.02; 0.01) | 0.00 (0.00; 0.01) |
| (b) Among populations |  |  |  |  |
| Body size | 0.22 (0.12; 0.36) | 0.37 (-0.09; 0.74) | 0.19 (-0.38; 0.63) | 0.12 (-0.73; 0.83) |
| Mesh width | 0.05 (-0.01; 0.12) | 0.08 (0.02; 0.17) | 0.22 (-0.53; 0.72) | 0.08 (-0.76; 0.79) |
| Web surface | 0.02 (-0.03; 0.09) | 0.02 (-0.02; 0.08) | 0.06 (0.00; 0.14) | 0.04 (-0.75; 0.77) |
| Egg load | 0.00 (-0.01; 0.01) | 0.00 (-0.01; 0.01) | 0.00 (-0.00; 0.01) | 0.00 (0.00; 0.00) |
| (c) Within populations (residual) |  |  |  |  |
| Body size | 0.71 (0.63; 0.79) | **0.33 (0.26; 0.41)** | **0.33 (0.26; 0.41)** | 0.14 (-0.03; 0.30) |
| Mesh width | **0.24 (0.18; 0.31)** | 0.75 (0.67; 0.84) | **0.56 (0.50; 0.61)** | 0.09 (-0.08; 0.25) |
| Web surface | **0.24 (0.18; 0.30)** | **0.40 (0.34; 0.48)** | 0.71 (0.63; 0.80) | 0.11 (-0.02; 0.24) |
| Egg load | 0.02 (-0.00; 0.05) | 0.01 (-0.01; 0.04) | 0.02 (-0.00; 0.04) | 0.03 (0.03; 0.04) |

## Discussion

Using a standardized hierarchical sampling design and multivariate modelling, we found *Araneus diadematus* to decrease in size with increasing levels of local urbanization, and independently, to adjust different components of its web-building behaviour. These changes appear to be adaptive (i.e. beneficial) and follow predictions of prey availability reduction and the urban heat island effect in response to urbanization. The putative adaptive value of these behavioural changes is supported by the fact that *A. diadematus* is able to maintain consistent population densities across broad gradients of urbanization, when many orb web spiders in the same communities cannot (Dahirel et al., 2017). Interestingly, while web building behaviours are intrinsically connected to body size, their responses to urbanization are here decoupled.

*Araneus diadematus* experienced size reduction in response to local urbanization only. The temperature-size rule, found in many arthropods, predicts that due to increased metabolic rates and costs, individuals are smaller at higher temperatures (Atkinson, 1994; Horne et al., 2015). As the spatial scale of urban heating matches the spatial scale at which size reduction is observed (Kaiser et al., 2016; Merckx et al., 2018), the Urban Heat Island Effect provides a valid explanation for the observed patterns (Merckx 2018, but see Entling, Schmidt-Entling, Bacher, Brandl, & Nentwig, 2010; Puzin, Leroy, & Pétillon, 2014). However, the impact of small temperature changes on spider body size remains to be studied, and resource availability during development is a well-documented influence and a potentially stronger driver of spider adult size (DiRienzo & Montiglio, 2016; Kralj-Fišer et al., 2014; Mayntz et al., 2003). Higher prey abundance in urbanized spots of the Sydney area was for instance associated with a larger body size in the spider *Nephila plumipes* (Lowe et al., 2014; Lowe, Wilder, & Hochuli, 2016). In our study region, prey size and prey biomass availability decreased with urbanization (Dahirel et al., 2017). The among-studies discrepancy in spider body size shifts cannot be easily reconciled with a major role of the Urban Heat Island effect, but is fully consistent with a prominent role of resource availability. Interestingly, we did not detect a spider size reduction in response to landscape-level urbanization (Table 2, Fig. 2), even though prey body size declines equally in response to urbanization at both scales (Dahirel et al., 2017). Several urban environmental changes, such as temperature increases or limited resource availability may be playing a role, their effects adding at the local scale only. Pollutants such as heavy metals also have a negative impact on body size (Ramirez et al., 2011), but we expect their influence here to be limited, as pesticide and heavy metal levels appear unrelated to urbanization in our study area (Merckx et al., 2018; T. Merckx, pers. comm.). Alternatively, resource loss may be the main mechanism driving size-reduction with urbanization, which is only observed at the local scale because spiders are able to alter their web-building behaviour to compensate for prey biomass loss at the landscape, but not at the local scale.

Changes in foraging behaviour in response to urbanization were scale-dependent: spiders built smaller webs with smaller mesh width in response to local urbanization, but increased their web surface with urbanization at the landscape scale (Fig. 2). Additionally, web surface responded to urbanization at both scales independently of mesh width variation (surface – mesh correlation ≈ 0; Table 3), meaning that web surface changes can tentatively be interpreted in terms of corresponding silk investment changes. Webs with smaller mesh are considered better at retaining prey (Blackledge & Zevenbergen, 2006), at the cost of a smaller web surface and therefore fewer prey intercepted for a given investment in silk production. When mesh width is held constant, as it is in response to landscape-level urbanization, increased silk investment leads to an increase in web surface (Figs 1, 2) and therefore an increase in the number of prey caught (Prokop & Grygláková, 2005; Venner & Casas, 2005). Because of their larger diameter, larger webs are also built with longer radial silk threads, meaning they can stop prey with higher kinetic energy (i.e. bigger prey) without breaking down (Harmer, Clausen, Wroe, & Madin, 2015). Although the importance of these large prey in particular for orb web evolution (Blackledge, 2011; Eberhard, 2013; Harmer et al., 2015) and individual performance (Harmer et al., 2015; Venner & Casas, 2005) remains controversial, selective advantages to having webs that maximize prey biomass in general are clear (Harmer et al., 2015). Temperature can also have a positive effect on silk production and hence web surface (Barghusen et al., 1997; Vollrath et al., 1997). However, the Urban Heat Island effect is stronger at the local, rather than landscape scale in our study region (Kaiser et al., 2016; Merckx et al., 2018), and is too weak (about 1-2°C) to explain the observed increase in web surface (in *Araneus diadematus*, a 12°C increase was needed to obtain surface changes similar to those we observed; Vollrath et al., 1997). Prey size (and thus biomass) decreased with urbanization at both scales (Dahirel et al., 2017). In this context, both the reduction of mesh size at local scales and the increase in capture area at larger spatial scales can be seen as adaptive responses to urbanization, respectively increasing prey retention and prey capture efficiency. These adaptations potentially contribute to the persistence of *Araneus diadematus* across urbanization gradients, and confirm that biotic interactions can be important drivers of phenotypic changes in urban environments (Alberti et al., 2017).

Web-building in orb-weaving spiders has previously been shown to be both size-constrained (Bonte et al., 2008; Gregorič et al., 2015), but also highly variable depending on the environment (Herberstein & Tso, 2011); the net effect in the case of environmental changes influencing body size was so far unknown. We show that correlations among our studied traits are only present within populations (residual covariance matrix) and not at the among-population and among-landscape levels (Table 3). These absences of correlation must be interpreted with some caution, as our credible intervals are wide. Nonetheless, our results suggest that while body size constrains web-building strategy within populations (Gregorič et al., 2015), it does not substantially limit the possibilities of spiders to alter their web-building behaviour in response to urbanization. The observed web building flexibility is thus a direct response to urban selection pressures, and not an indirect consequence of body size shifts. Additionally, these results confirm that the existence of trait correlations/syndromes in one environmental context or at one biological organisation level may not be informative with respect to their maintenance across contexts/ levels (e.g. Fischer, Ghalambor, & Hoke, 2016; Peiman & Robinson, 2017).

Changes in web-building behaviour in relation to urbanization may both originate from plasticity and/or evolutionary changes (Herberstein & Tso, 2011). Disentangling the relative contribution of both is however difficult in observational studies (Merilä & Hendry, 2014). We take advantage of our two-spatial scales design, and the fact that plasticity and genetic adaptation are generally expected to appear in response to finer-grained and coarser-grained environmental variation, respectively (Richardson et al., 2014), to emit hypotheses on the relative contributions of these two mechanisms. Mesh width varies with urbanization at the local but not at the landscape scale. This would indicate a more important role of plastic responses, in accordance with experimental work demonstrating high within-individual behavioural flexibility in response to food availability and prey spectrum (reviewed by Herberstein & Tso, 2011). By contrast, silk investment, reflected in web surface, adaptively increased in response to landscape-scale urbanization. With the caveat that some drivers of web building variation may have been overlooked, this suggests here a putative role of genetic adaptation in urbanization-driven silk production changes. We acknowledge our results only provide first hints in this direction, and that further research will be needed to quantify the contribution of genetic selection relative to developmental and behavioural plasticity.

Costs associated with urban life were especially prominent at the local scale. Indeed, at this scale, body sizes and web capture surfaces were smaller, and lower fecundity was observed despite stable population densities (Fig. 2, Table 2). Egg load did not, however, covary with body size or web traits at either the inter- or intra-population levels. Reduced fecundity is therefore unlikely to be a direct consequence of altered web-building behaviour or a reduced body size, but rather an additive response to environmental changes associated with urbanization. Food limitation has strong negative impacts on spider lifetime fecundity even in the absence of survival costs (Kleinteich, Wilder, & Schneider, 2015) or body size changes (Miyashita, 1990). The lower fecundities observed in locally urbanized sites may therefore indicate that spiders were unable to fully compensate for reduced prey biomass at this scale despite shifts in web-building, and despite smaller size leading to reduced requirements. Other –here unmeasured-stressors related to urbanization may also lead to reduced fecundity and capture area in locally urbanized sites: heat waves (Kingsolver, Diamond, & Buckley, 2013), pollution (fecundity costs: Hendrickx, Maelfait, Speelmans, & Straalen, 2003; web surface costs: Benamú et al., 2013; but see Ramirez et al., 2011), or repeated web destructions and rebuilding following human disturbance (Tew, Adamson, & Hesselberg, 2015). Overall, *Araneus diadematus* was comparatively much better at dealing with landscape-scale urbanization; indeed, both fecundity and population densities remained unaffected by urbanization at this scale. This suggests that traits that responded at this scale (namely web surface) are more important to prey capture success and biomass gain than the others (mesh width) (Blackledge & Eliason, 2007). Conversely, benefits, costs and trade-offs associated with increased silk production may be detectable in other life-history dimensions than the ones we explored in the present study, such as development time and adult longevity (both expected to increase with resource restriction; Kleinteich et al., 2015; Kralj-Fišer et al., 2014), or dispersal, which may be highly risky in urban fragmented environments for passively dispersing organisms (Cheptou, Carrue, Rouifed, & Cantarel, 2008).

From an applied perspective, our results confirm the oft-stated importance of maintaining local greenspots in highly urbanized landscapes (e.g. Philpott et al., 2013), as even generalist “winning” species such as *A. diadematus* may benefit from them. Additionally, even though spider abundance stayed constant, webs were up to 20% smaller in locally urbanized sites. Spiders are important predators in natural and anthropogenic landscapes (Birkhofer, Entling, & Lubin, 2013; Foelix, 2010): our results highlight the necessity of accounting for changes in functional, and not only numerical predator responses to accurately quantify ecosystem services provided by green infrastructures in urban environments. Further comparisons among successful species that differ in their silk production strategy (e.g. *A. diadematus*, which destroys and recreates webs regularly, versus *Nephila plumipes*, which build semi-permanent webs; Lowe et al., 2014) may highlight commonalities and help shed further light on the costs/ benefits balance of adaptation to urban life.

## Authors’ contributions

DB conceived the study and designed methodology; MDC and PV collected the data; MD and MDC analysed the data; MD led the writing of the manuscript. All authors contributed critically to the draft and gave final approval for publication.

## Acknowledgments

We are grateful to Jasper Dierick for his assistance during field sampling, and to Hans Matheve for GIS calculations and site selection. We thank Niels Dingemanse and three anonymous reviewers for their comments on a previous version of this paper. This article is part of the Belspo-funded IAP project P7/04 SPEEDY (SPatial and environmental determinants of Eco-Evolutionary DYnamics: anthropogenic environments as a model). MD is funded by a postdoctoral grant from the Fyssen Foundation.

## Data accessibility

Data will be made available on Dryad upon final article acceptance.

