## Supplementary Materials for "Urbanization-driven changes in web-building are decoupled from body size in an orb-web spider"

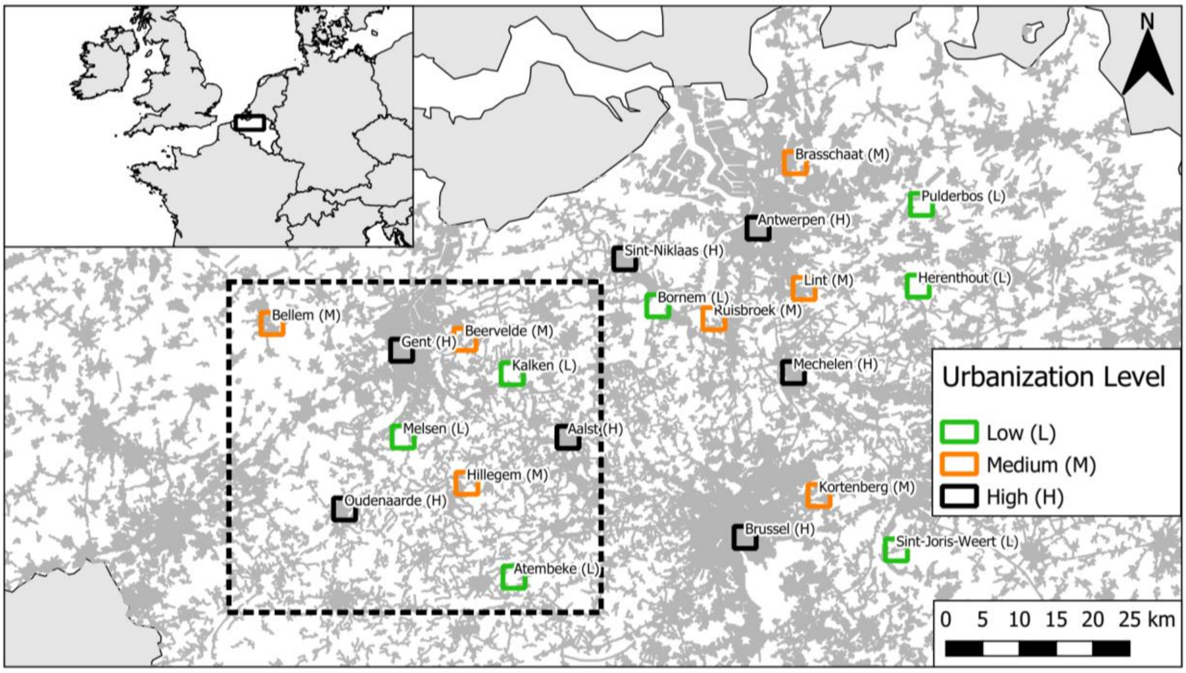


**Supplementary Figure 1** Location of the 21 studied plots within northern Belgium. Plot colour denotes landscape-scale urbanization level. Three 200 × 200 m sites were sampled per plot. Dark grey zones correspond to artificialized areas after CORINE Land Cover (European Environmental Agency, 2016). Information on prey availability in the previous study by Dahirel *et al.* (2017) was collected in the nine plots within the dotted line rectangle. Inset: Location of the study area within the broader western Europe.

Supplementary Material for: **Maxime Dahirel, Maarten De Cock, Pieter Vantieghem and Dries Bonte.** Urbanization-driven changes in web-building are decoupled from body size in an orb-web spider

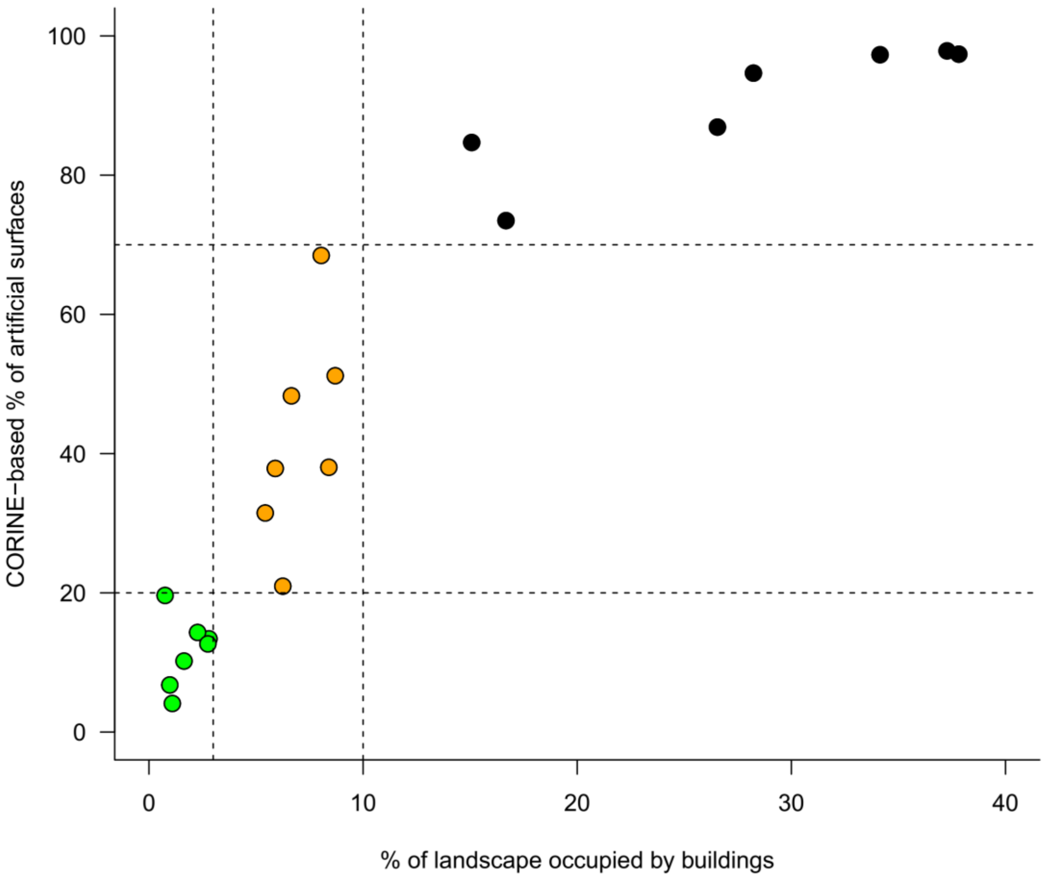


**Supplementary Figure 2** Comparison between landscape-level urbanization values based on the percentage of buildings and based on the percentage of artificialized surfaces *sensu* CORINE Land Cover (European Environmental Agency, 2016; excluding urbanized greenspots). Spearman correlation between the two metrics: 0.94 ($N$ = 21 landscapes, $p = 5.17 \times 10^{-6}$). Colours refer to urbanization levels as used in the article (green: low urbanization, orange: moderate urbanization, black: high urbanization), with dotted lines corresponding to the thresholds used to separate these levels.
